## Supplementary figures and images for "ZFT is the major iron and zinc transporter in *Toxoplasma gondii*"

### Supplemental Figures

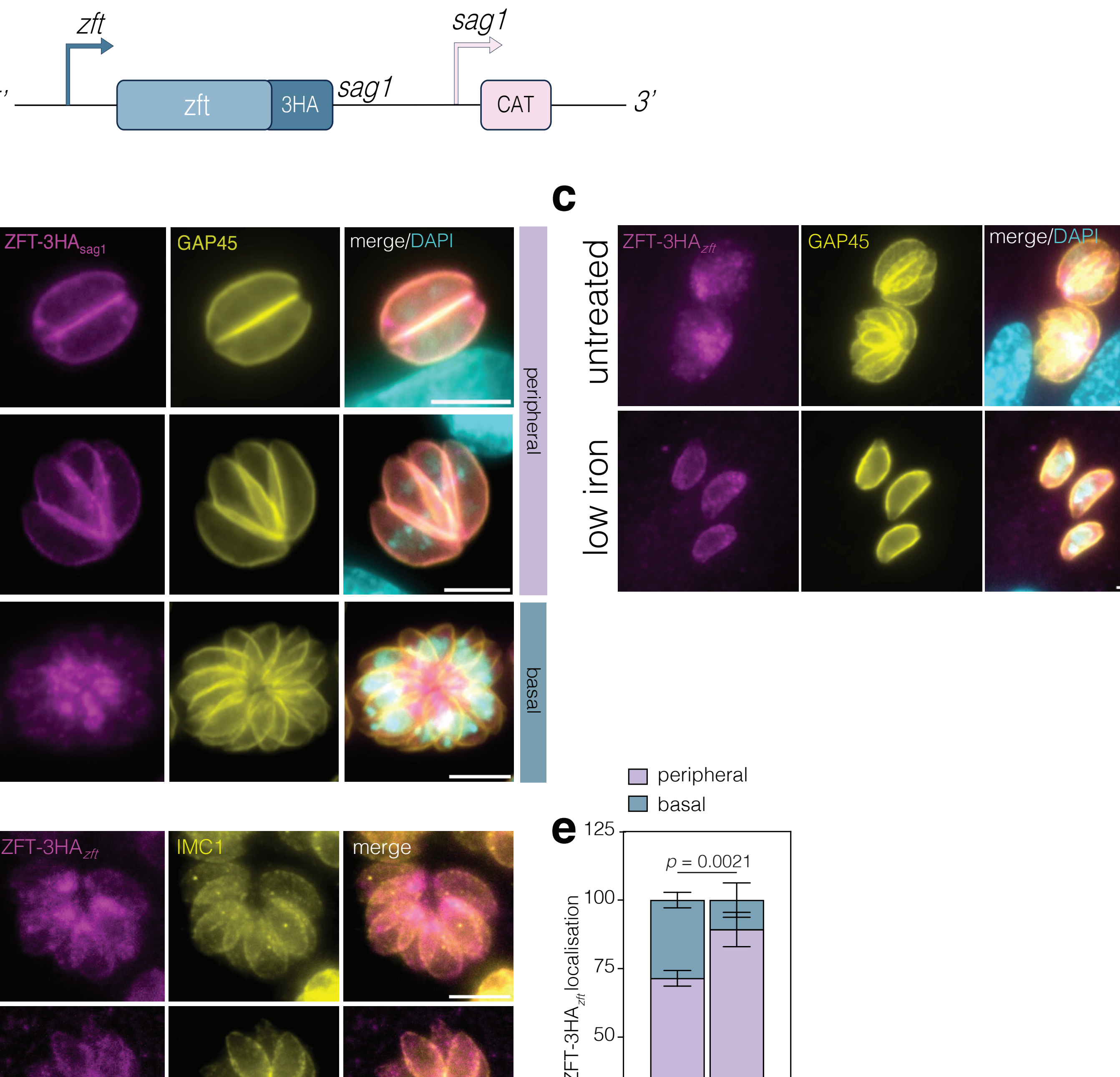

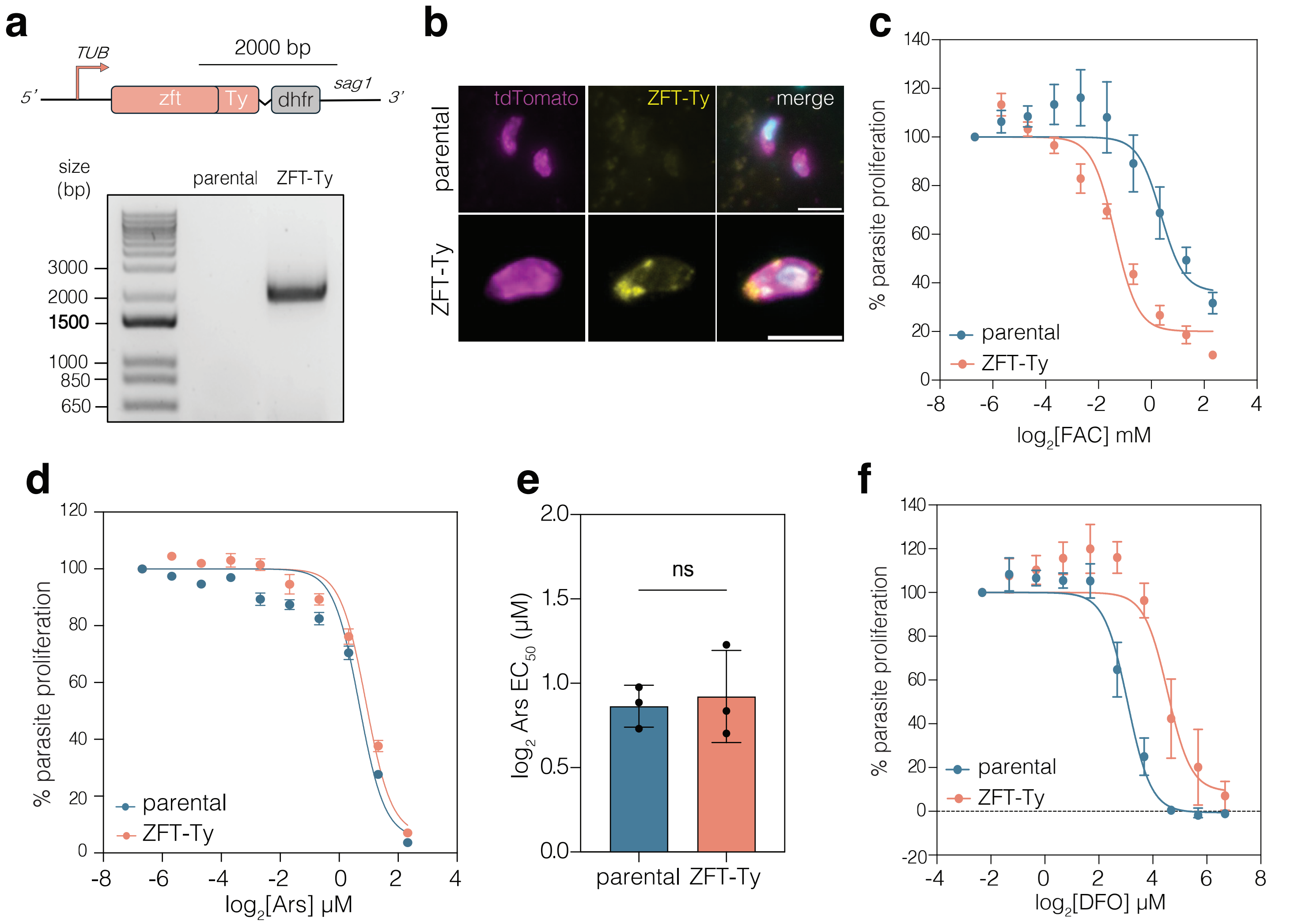

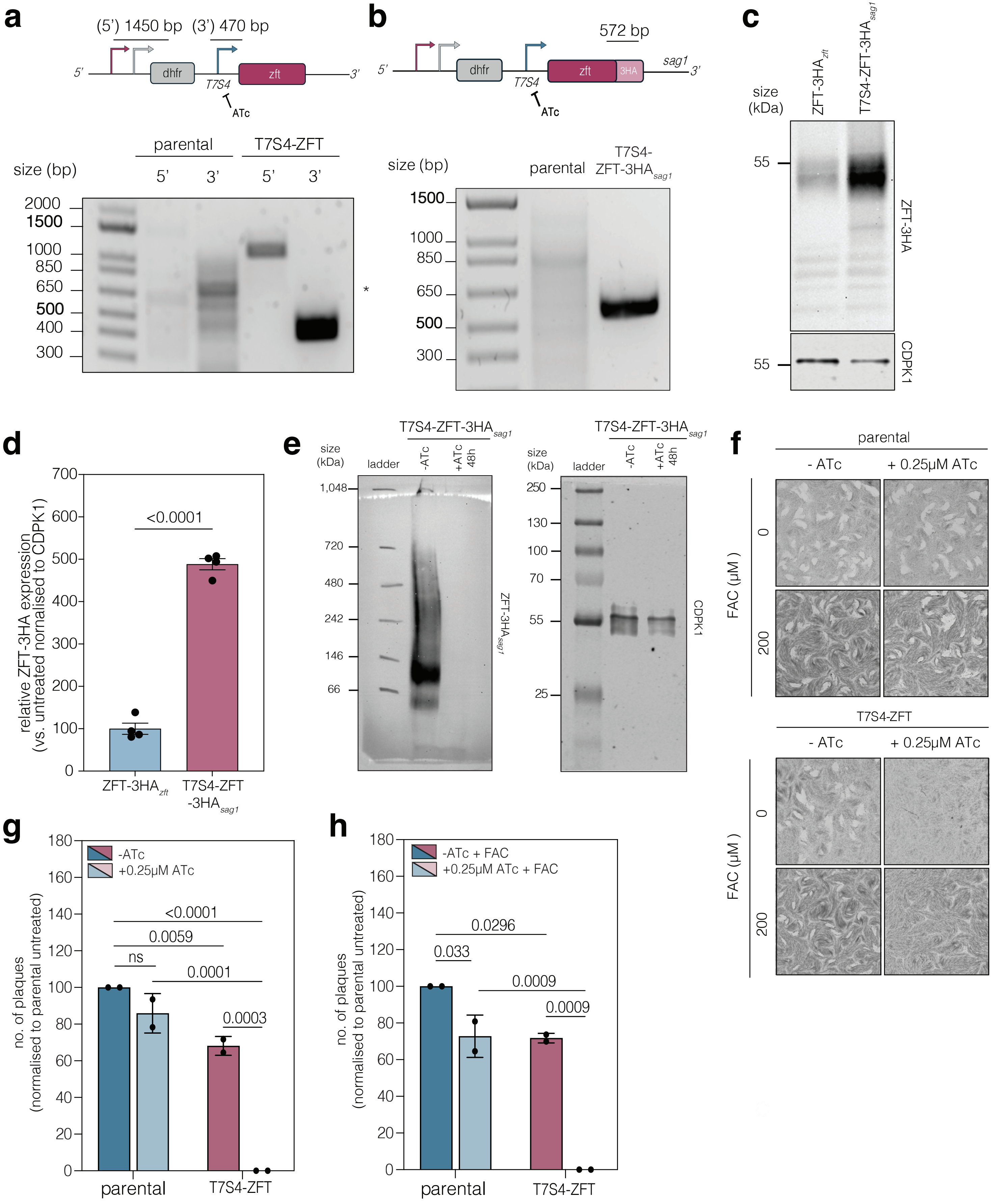

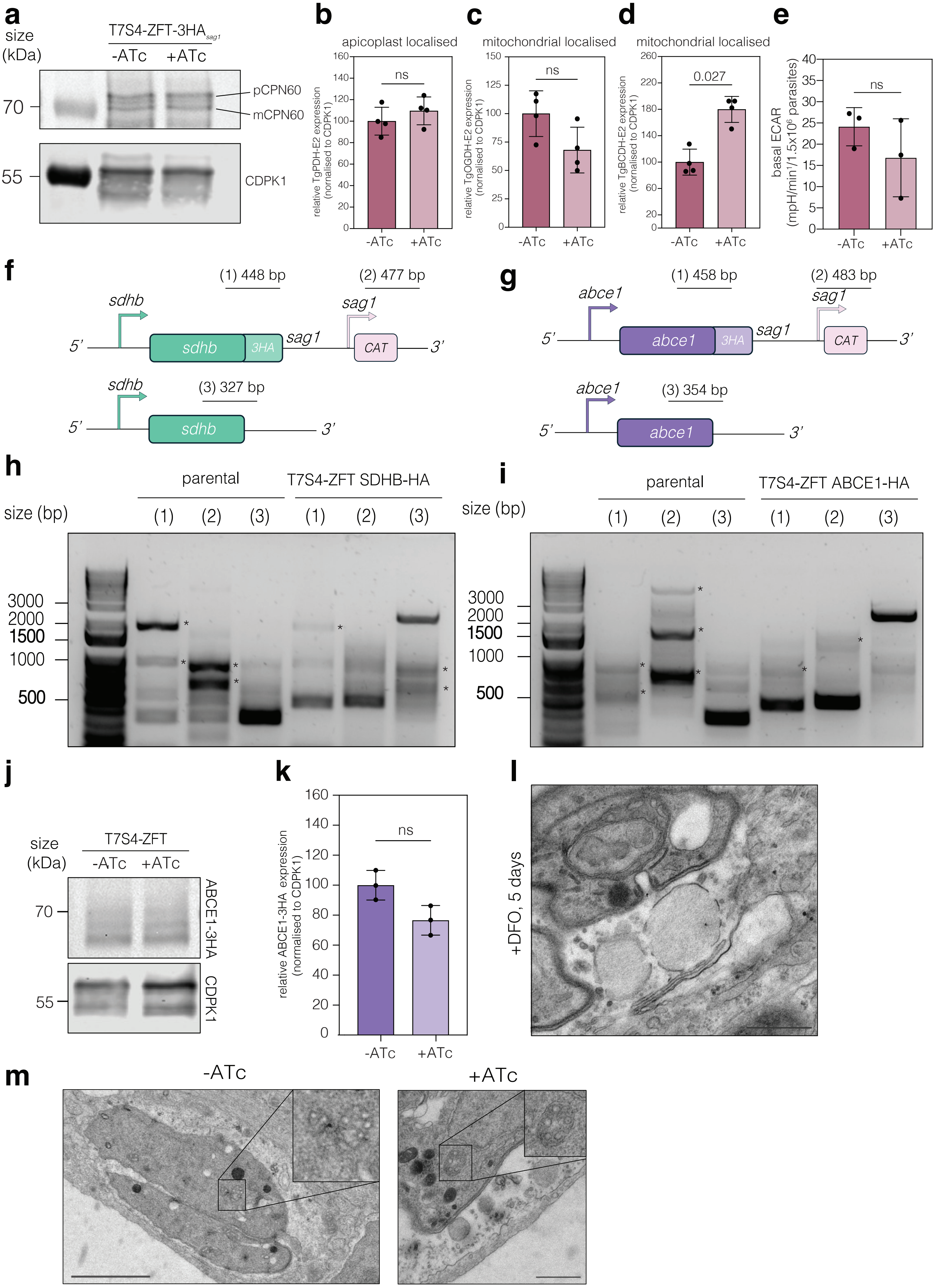

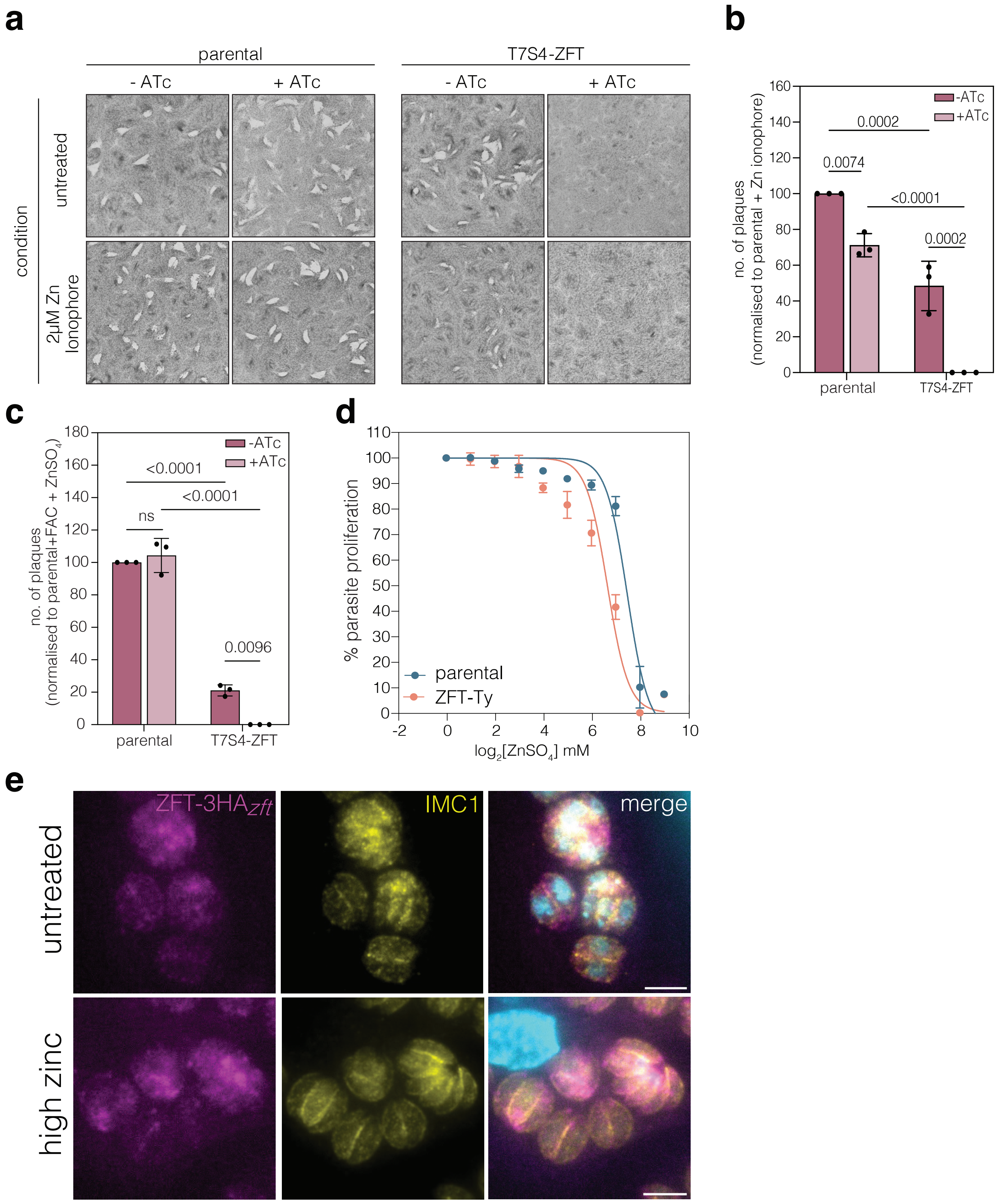

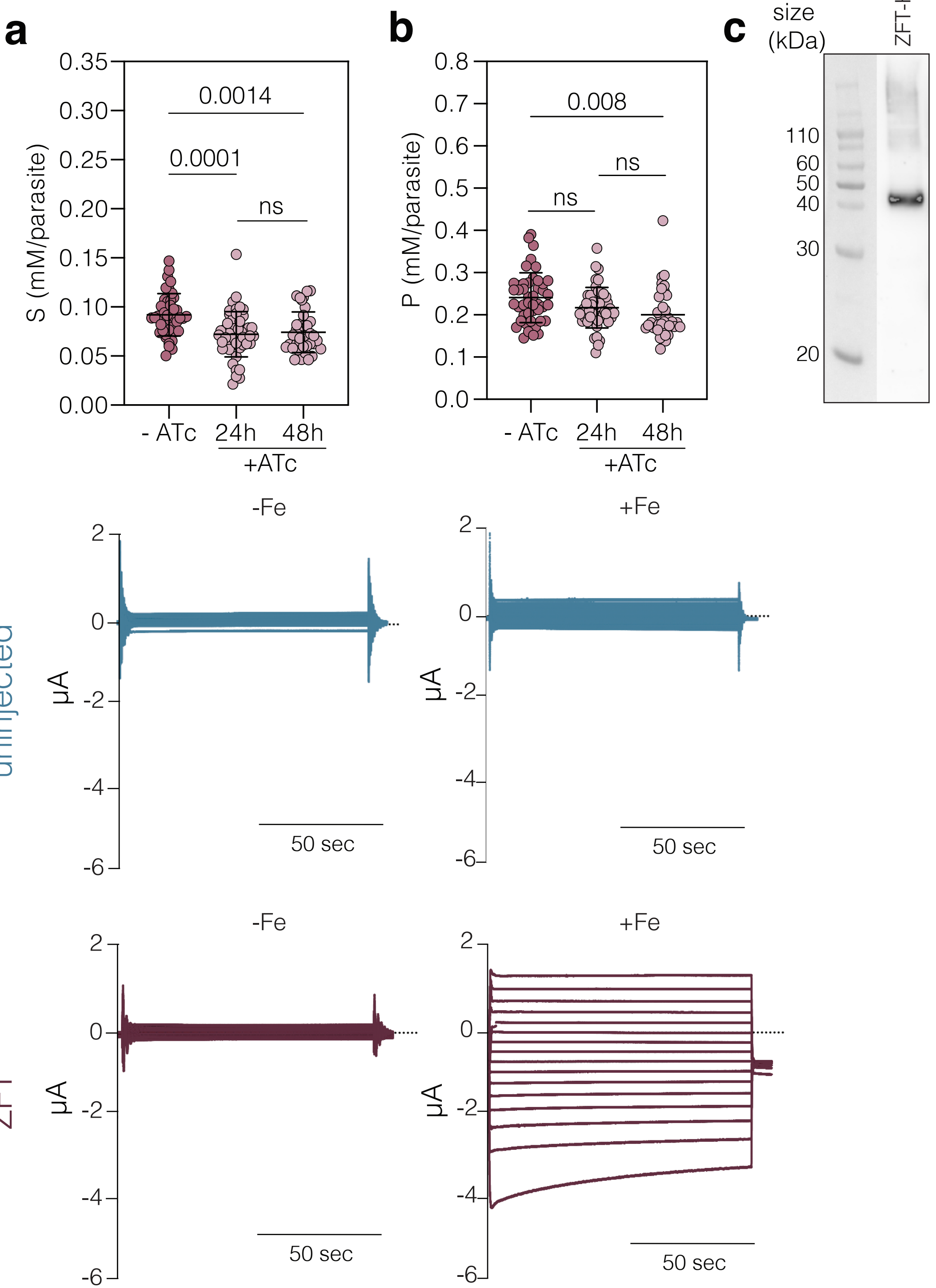
